# SMARTcleaner: identify and clean off-target signals in SMART ChIP-seq analysis

**DOI:** 10.1101/269365

**Authors:** Dejian Zhao, Deyou Zheng

**Author notes:** Correspondence should be addressed to, Deyou Zheng, Ph.D., 1217.

## Abstract

**Background:** Noises and artifacts may arise in several steps of the next-generation sequencing (NGS) process. Recently, a NGS library preparation method called SMART, or <u>*S*</u>witching <u>*M*</u>echanism <u>*A*</u>t the 5’ end of the <u>*R*</u>NA <u>*T*</u>ranscript, is introduced to prepare ChIP-seq (chromatin immunoprecipitation and deep sequencing) libraries from small amount of DNA material. The protocol adds Ts to the 3’ end of DNA templates, which is subsequently recognized and used by SMART poly(dA) primers for reverse transcription and then addition of PCR primers and sequencing adapters. The poly(dA) primers, however, can anneal to poly(T) sequences in a genome and amplify DNA fragments that are not enriched in the immunoprecipitated DNA templates. This off-target amplification results in false signals in the ChIP-seq data.

**Results:** Here, we show that the off-target ChIP-seq reads derived from false amplification of poly(T/A) genomic sequences have unique and strand-specific features. Accordingly, we develop a tool (called “SMARTcleaner”) that can exploit the features to remove SMART ChIP-seq artifacts. Application of SMARTcleaner to several SMART ChIP-seq datasets demonstrates that it can remove reads from off-target amplification effectively, leading to improved ChIP-seq peaks and results.

**Conclusions:** SMARTcleaner could identify and clean the false signals in SMART-based ChIP-seq libraries, leading to improvement in peak calling, and downstream data analysis and interpretation.

## Background

In the past decade, deep sequencing by next generation sequencing (NGS) has been widely applied in nearly all fields of biological research, in which information from biological processes (e.g., transcription and protein-DNA interaction) can be converted to DNAs for sequencing [1-4]. NGS is a complex procedure involving DNA/RNA isolation, library preparation, deep sequencing, data processing and interpretation. Each of these steps can introduce biases and artifacts, but the first step - preparation of NGS libraries is arguably the most critical phase as errors can be propagated to later steps, if not carefully controlled [5, 6]. Among them, PCR amplification is a major source of bias due to the fact that not all fragments are amplified with the same efficiency [5].

As powerful as NGS technology is, its application with limited amounts of biological material, for example, DNA or RNA isolated from a very small number of cells, remains a challenge. This is primarily due to the low efficiency in ligating targeted DNA/RNA fragments to the NGS sequencing adaptors, leading to a drop of sequencing reads for low copy DNA/RNA molecules present in a sample [7]. In addition, ligation requires double-stranded DNA (dsDNA) inputs and may result in cross- and self-ligation adaptor byproducts [8]. To overcome these limitations, SMART, a template switching method, was developed and used initially for transcriptome analyses, such as CAGE, RNA-seq (including small RNA-seq), and single-cell RNA-seq [9-12]. By using single-step adapter addition, the SMART technology achieves a much-needed sensitivity to accurately amplify picogram quantities of nucleic acids.

The SMART method was adapted for preparing NGS libraries from DNA templates in 2014 by tailing an adaptor to the 3’ end of a target DNA sequence and later amplifying the sequence by template switching. This modification allows quick preparation of DNA libraries from picogram quantities of DNA molecules [7]. Soon, this strategy was applied to ChIP-seq studies with human, mouse and yeast samples [13-19], and it is one of the few currently available protocols for ChIP-seq studies of small cell numbers [20, 21]. Here, a stretch of Ts is added to DNA templates in the tailing step, which is subsequently hybridized to a poly(dA) primer used to copy DNA (**Fig. 1a**). It is conceivable that the poly(dA) primer, however, can lead to signals amplified from non-targeted genomic regions containing consecutive Ts. Indeed, a recent study of SMART ChIP-seq reads revealed a strong bias of base constitution at the 3’ end of the sequenced reads that are enriched near long (≥12bp) poly(T/A) containing genomic loci [14]. The authors proposed a computational strategy to reduce this bias by normalizing the ChIP-seq data for the genomic abundance of different polyN tracts, but only achieved partial success [14]. Here, we revisited this problem and demonstrated that the unique features of the falsely amplified reads can be exploited to effectively remove artifact ChIP-seq reads from SMART protocols. We implemented this idea in the software SMARTcleaner. Testing multiple published ChIP-seq data, we showed that SMARTcleaner could properly identify and remove artifact reads in both paired-end (PE) and single-end (SE) ChIP-seq data, leading to improved ChIP-seq results.

**Fig. 1.**
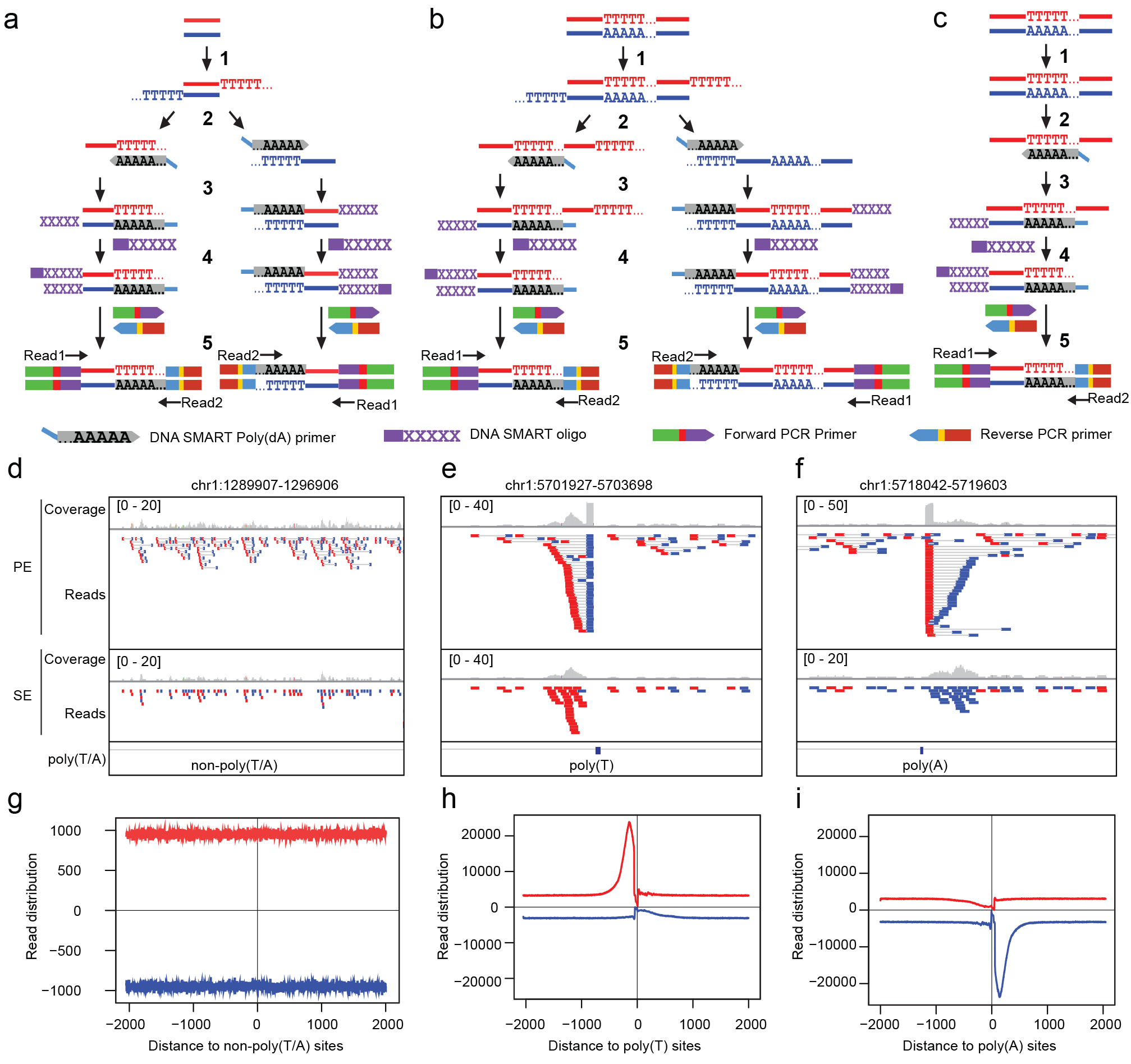
Strand-specific amplification of non-targeted sequences at poly(T/A) sites in the SMART ChIP-seq analysis. **a.** Flowchart of the SMART ChIP-seq procedure at non-poly(T/A) sites, adapted from the user manual of the kit (http://www.clontech.com/xxclt_ibcGetAttachment.jsp?cltemld=99449). **b,c**. Modified flowcharts to show annealing of the SMART poly(dA) primers to nontailed Ts within targeted (**b**) or non-targeted (**c**) DNA templates, leading to strand-specific amplification at poly(T) sites. For poly(A) sites, false amplification occurs to the opposite strand. **d-f**. ChIP-seq read densities at three randomly picked non-poly(T/A) and poly(T/A) sites. The data is from SRR3229031 (Additional file 1: Table S1, Dataset 1), and lntegrative Genomics Viewer (lGV) [32] is used to show the ChIP-seq reads from paired-end (PE) or single-end (SE) sequencing. For PE, read1 and read2 are shown as pairs, with reads mapped to “+” and “−“ strands in red and blue, respectively. For SE, only Read1 (extracted from PE data) is shown. **g-i**. Aggregated read distribution at non-poly(T/A) and poly(T/A) sites. ln h and i, poly(T/A) sites were defined as those with ≥ 12 consecutive T or A in the human reference genome. To define non-poly(T/A) sites, we first selected genomic regions that are > 4 kb in length and > 1 kb away from poly(T/A) sites, and then take the 2kb regions around the middle points. ln total, we got 301,474 non-poly(T/A) sites, 338,568 poly(T) sites, and 336,703 poly(A) sites. Refer to the Method section (SE mode, Additional file 2: Figure S6) for the calculation of read distribution.

## Results

### Strand-specific false priming and amplification at the poly(T/A) sites

When the SMART protocol (or kit) is applied to prepare NGS libraries from DNA fragments, such as those from chromatin immunoprecipitation (IP), there are five steps, 1) 3’ T-tailing, 2) annealing of DNA SMART poly(dA) primer to the T-tails, 3) primer extension by the SMARTScribe^TM^ reverse transcriptase (RT), 4) template switching and extension by RT using SMART oligo, and 5) PCR-mediated addition of Illumina adapters and subsequent amplification (**Fig. 1a**). As mentioned previously [14], the SMART poly(dA) primers can anneal to poly(T) sequences that are either located within the IP-DNA fragments (**Fig. 1b**) or present in non-target DNA fragments (i.e., the DNA fragments pulled down during IP non-specifically) (**Fig. 1c**). In both cases, the Ts are from genomic sequences and are not added during the T-tailing process. After amplification, sequencing, and read mapping (note that only one strand of the dsDNA is sequenced), ChIP-seq reads from poly(T/A) genomic DNAs, due to false priming and amplification, will accumulate next to the poly(T/A) sites in a clear strand-specific manner because the poly(dA) primers only anneal to the DNA strand containing poly(T). To illustrate this, we examined the reads in a human ChIP-seq sample (Additional file 1: Table S1, Dataset 1, SRR3229031) that was prepared using the Clontech DNA SMART ChIP-seq kit and by PE sequencing [14]. As this particular dataset was obtained from sequencing of control samples (i.e., input DNA), no genomic regions would be expected to show ChIP-seq read enrichment. lndeed, at non-poly(T/A) sites, we did not find accumulations of reads on either “+” or “−“ strands (**Fig. 1d**). However, at poly(T/A) sites, we observed that the Read2 of the PE reads were piled up either at the upstream of the poly(T) sites (with respect to the reference “+” strand) (**Fig. 1e**) or at the downstream of the poly(A) sites (**Fig. 1f**), as reported [14]. If SE sequencing had been performed, the accumulation of reads would still be observed, but the precise location information provided by Read2 would not be available (**Fig. 1e,f**), because only Read1 (**Fig. 1a-c**) would be sequenced. Genome-wide analysis of read distribution aggregated over poly(T/A) sites further illustrate these patterns (**Fig. 1g-i**). The width of the peaks indicates the range where the false fragments are located near the poly(T/A) sites (**Fig. 1g-i**).

### Random false priming and amplification at consecutive and intermittent poly(T/A) sites

We reasoned that the SMART poly(dA) primers can anneal to and amplify poly(T) sequences, allowing some degree of mismatch. The PE sequencing data in the SRR3229031 dataset allowed us to identify exactly the ChIP-seq fragments that were artifacts from the poly(T/A) genomic sites, because the Read2 of the fragments would be piled up at the end of poly(T/A) (**Fig. 1e,f;** Additional file 2: Figure S1). We should point out that the second reads of the PE sequences submitted to the SRA database have been cut by 10 bp from the 3’ end by the authors [14], resulting in a 10 bp gap between the poly(T/A) sites and the end of the Read2 (Additional file 2: Figure S1).

We counted the numbers of ChIP-seq Read2 that mapped to the 9,698,838 poly(A) and 9,796,521 poly(T) sequences containing a minimal of five consecutive As or Ts, respectively, in the human genome (hg38). Like a previous study [14], we found that the median counts for the regions with 5 to 11 consecutive A or T were 1, while the median for regions with 12 As or Ts was doubled, indicating that the false priming event occurs primarily at sites with 12 or more consecutive poly(T/A) bases (Additional file 2: Figure S2a; Wilcoxon test, *p*-value < 2.2e-16). Nevertheless, there were large variations at the poly(T/A) sites of the same length, a common phenomenon due to the randomness in primer annealing and sequencing (Additional file 2: Figure S2a). To consider mismatching during priming, we focused on short poly(T/A) sites (≤8bp) that by themselves cannot be efficiently used for false priming but jointly may be. We found that read numbers mapped to two such sequences disrupted by one mismatch nucleotide were significantly reduced, compared to those without disruption, indicating reduced efficiency of false priming (Additional file 2: Figure S1c,d, Figure S2b). Moreover, an insertion of two or three mismatch nucleotides basically abolished false priming (Additional file 2: Figure S2b). In short, our analysis confirmed that false priming occurs significantly at regions containing a consecutive sequence of ≥12 As or Ts and the resultant artifact reads should be excluded from ChIP-seq data analysis.

### SMARTcleaner: identification and cleaning of falsely primed fragments

Based on the above information of the false priming event in SMART ChIP-seq studies (**Fig. 1**, Additional file 2: Figure S2), we developed a computational tool, SMARTcleaner, to remove the ChIP-seq artifact signals. It has two modes (PE mode and SE mode) to accommodate the two sequencing options during ChIP-seq. In PE mode, a genome (FASTA) sequence file and ChIP-seq read alignment files (in bam format) are taken as input, and “cleaned” bam files are generated with the reads predicted from false priming removed and saved in the “noise” bam files. In SE mode, it takes a list of consecutive and interrupted poly(T/A) genomic sites (Additional file 2: Figure S2), and bam files, and outputs cleaned bam files and noise bam files. The software is publicly available through github (https://github.com/dzhaobio/SMARTcleaner).

In PE mode, our tool removes ChIP-seq read pairs whose second reads mapped to poly(T/A) (see Methods). Analysis of pileup reads at individual poly(A/T) sites (**Fig. 2a,b**) and total read counts across all poly(A/T) sites (**Fig. 2c,d**) demonstrated clearly that reads from false priming in the SRR3229031 dataset were effectively identified and successfully removed by SMARTcleaner. Furthermore, applying the SMARTcleaner to ChIP-seq data from libraries constructed using a ligation method [14], we found that < 0.002% of PE reads were mistakenly removed, indicating that the PE mode is highly accurate. By comparison, artifact reads in the SMART-based data could be successfully removed, while their percentages (11-20%) varied among the different DNA shearing methods used for fragmentation (**Fig. 2e**). In addition, for the SMART-based data, the ChIP-seq fragment sizes calculated from the noise bam files were 21-43 bp shorter on average than those in the clean bam files, as expected, since the genomic poly(T/A) sequences were within ChIP fragments while tailed Ts were added to the ends of ChIP fragments. This observation is consistent with previous finding [14].

**Fig. 2.**
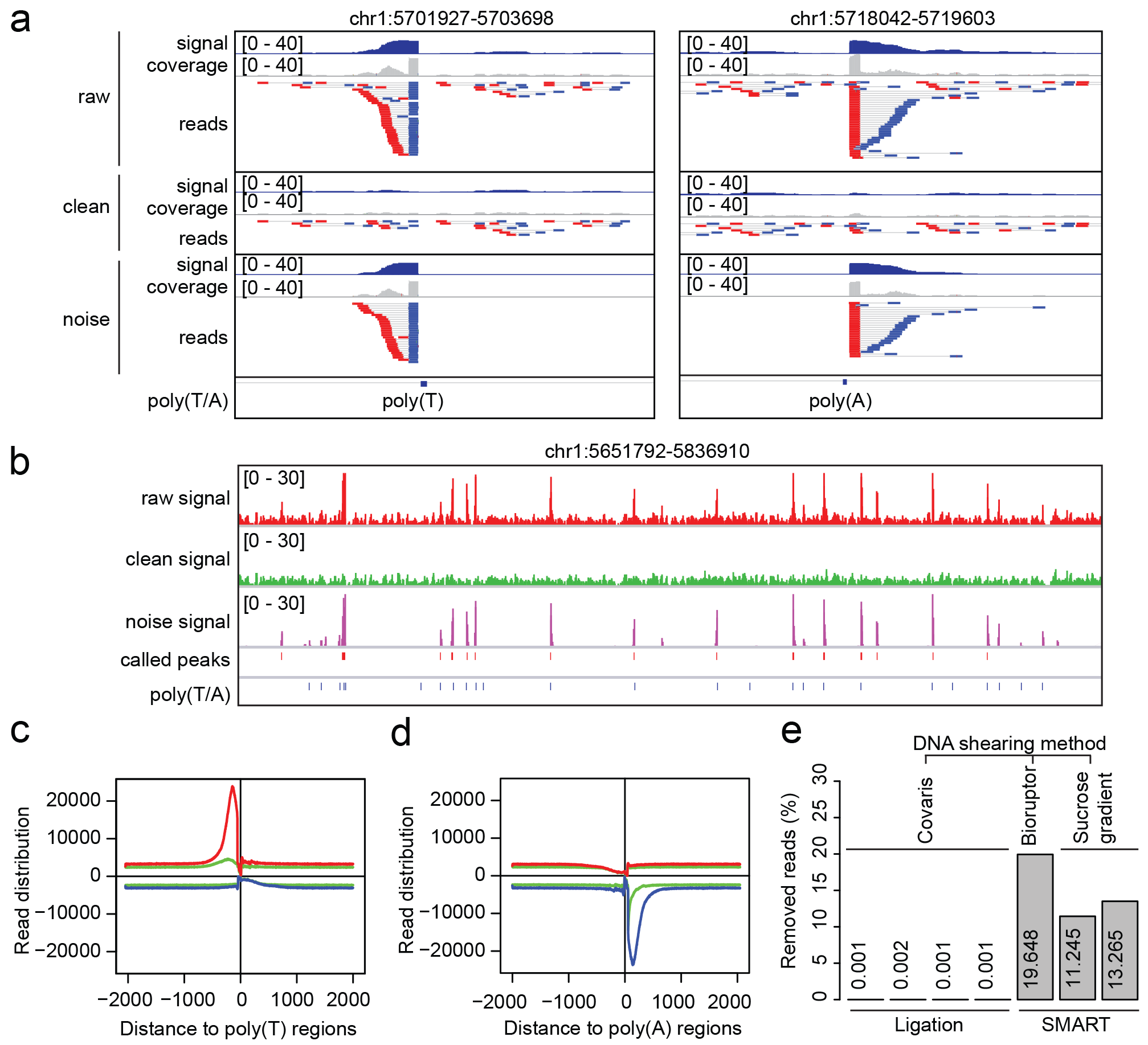
SMARTcleaner in PE mode. **a.** PE reads mapped to a poly(T) and a poly(A) locus before (raw) and after cleaning. **b**. A genomic region showing the read densities before and after cleaning. The “called peaks” refer to pre-cleaning peaks called using MACS2. **c,d**. Genome-wide read distribution at poly(T/A) sites before (red and blue lines) and after (green lines) cleaning. **e**. Percentages of removed reads at poly(T/A) sites in each sample. The samples from left to right are SRR3229030, SRR3286889, SRR3286890, SRR3286891, SRR3229031, SRR3286910, and SRR3286911 (Additional file 1: Table S1, Dataset 1).

In SE mode, the SMARTcleaner identifies and removes artifact reads by comparing read distributions in the “+” and “−“ strands near individual poly(T/A) sites, because false priming leads to reads accumulated in only one of the two strands (**Fig. 1**). To demonstrate its performance, we treated the above PE ChIP-seq reads as SE reads, by analyzing the Read1 data only. Again, analysis of pileup reads at individual poly(T/A) sites (**Fig. 3a,b**) and read counts aggregated over genome wide poly(T/A) sites (**Fig. 3c,d**) demonstrated that most artifact reads were removed effectively. However, the SE mode appeared less robust than the PE mode, because it mistakenly removed ~0.8% of reads in the ligation-based ChIP-seq data (**Fig. 3e**). The percentages of reads that were removed by the SE mode for the SMART-based datasets were similar to those using the PE mode (**Fig. 3e**).

**Fig. 3.**
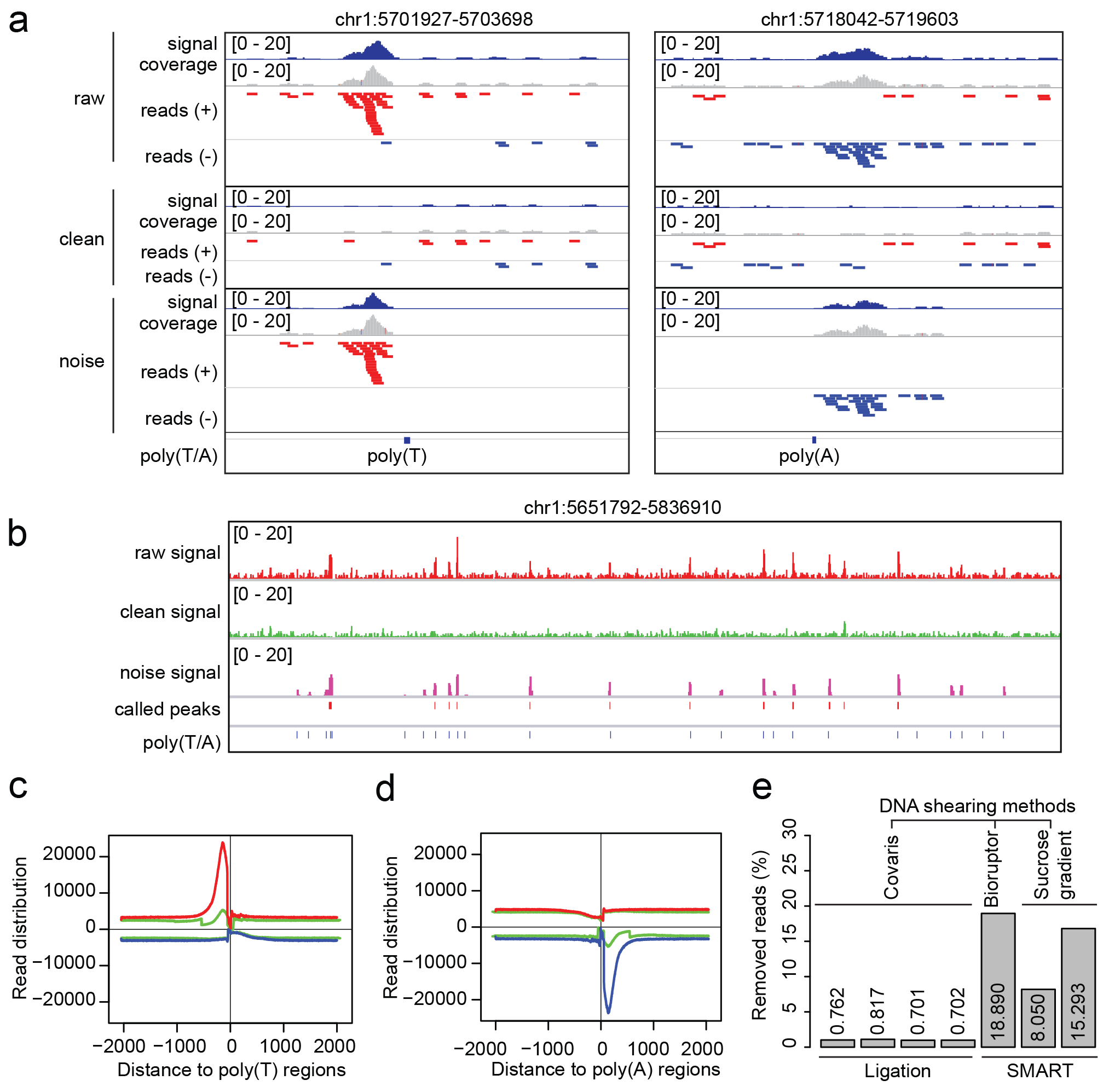
SMARTcleaner in SE mode. **a**. Two examples showing the cleaning results of SE mode at one poly(T) and one poly(A) locus. **b**. Cleaning result in a genomic region. **c,d**. Genome-wide reads distribution near the poly(T/A) sites before (red and blue lines) and after (green lines) cleaning. **e**. Percentages of removed reads at poly(T/A) sites in samples prepared by ligation or SMART protocols. The sample order is the same as in Fig. 2e.

In terms of computational efficiency, we tested both PE and SE modes on a PC (lntel(R) Xeon(R) CPU E5-2609 0 @ 2.40GHz, 32Gb memory, CentOS Linux release 7.3.1611). It took 30 min to clean 94 million reads in PE mode and 16 min to clean 47 million reads in SE mode, benchmarking with the SRR3229031 dataset. The PE mode requires more memory than the SE mode because the former reads the entire genome sequence into memory (for fast query) and keeps track of the end coordinates of Read2 at the genomic poly(T/A) sites.

### Evaluation of SMARTcleaner with published histone modification ChIP-seq datasets

To demonstrate the value of our tool and importance of removing artifact reads from false priming in the analysis of SMART ChIP-seq data, we first applied the SMARTcleaner to a public ChIP-seq dataset (Additional file 1: Table S1, Dataset 2) that studied H3K4me3 histone modification in HeLa cells using seven methods for preparing sequencing libraries from low-input IP DNAs, including SMART method [13]. The study also generated a PCR-free dataset as a gold standard reference, including three replicates using 100 ng DNA as starting material. For the other seven protocols, the starting material was either 1 ng or 0.1 ng, each with five replicates [13]. The original study was designed for comparing the performance of different ChIP-seq library preparation methods, but this dataset is ideal for evaluating our tool for three reasons. First, its gold standard data can be used for clearly evaluating artifacts introduced in PCR amplification. Second, the dataset is valuable for evaluating the effect of initial DNA inputs on false priming and amplification. Third, the known enrichment of H3K4me3 peaks at promoter regions [22] can be used as a metric to measure the impact of falsely called peaks.

In our test below, as a benchmark we chose the data from PCR-free method and Ascel2S method, which were consistently ranked at the top by multiple criteria in the original study [13]. Since the ChIP-seq libraries were sequenced by the single-end method, we applied SE mode to the alignment files, including control samples. Similar to the above finding in **Fig. 3e**, only a small percentage of ChIP-seq reads were removed by SMARTcleaner from the ligation-based datasets, 0.3% on average. For SMART-derived dataset, the average percentage was 3.0% for 1 ng and 5.3% for 0.1 ng starting DNA material (Additional file 2: Figure S3a). Next, we randomly sampled 6 millions of reads for each sample for calling H3K4me3 peaks using the software MACS2 [23], by the same criteria. We found that before read cleaning 12.1% and 17.1% of the H3K4me3 peaks, called from the 1 ng and 0.1 ng SMART protocols respectively, overlapped with poly(T/A) sites, but after cleaning the overlaps dropped to 6.2% and 8.1%, comparable to the numbers for PCR-free and Ascel2S samples (Additional file 2: Figure S3b). This result indicates that not all peaks in poly(T/A) sites are artifacts. The greater percentages of removed reads and peak overlaps with poly(T/A) sites for the 0.1 ng than the 1 ng dataset are consistent with the assumption of increased false priming when the input DNA material is lower, due to a reduced number of genuine target DNA templates. In addition, the percentages of H3K4me3 peaks mapping to promoters increased by 3.7% (1 ng) and 4.1% (0.1 ng) after cleaning reads in the SMART derived datasets, while the change (0.14%) is negligible for the PCR free and Ascel2S samples (Additional file 2: Figure S3c).

We also compared the SMART ChIP-seq peaks to the H3K4me3 peaks from PCR-free samples, using the peaks (n= 20,262) present in all three PCR-free datasets as the reference. The mean sensitivity (i.e., % PCR-free peaks detected in SMART) was 89.68% and 89.61% in pre- and post-cleaning samples (1ng DNA), indicating no difference in sensitivity. Same was observed for the samples using 0.1ng starting DNA material (Additional file 2: Figure S3d). However, the specificity (% SMART peaks found in PCR-free peaks) was increased from 89.25% to 90.42% for samples with 1ng DNA and from 87.11% to 89.85% for samples with 0.1ng DNA after cleaning the noise (Additional file 2: Figure S3e), indicating that the cleaning process improved the peak quality.

Next, we directly compared the pre- and post-cleaning H3K4me3 peak lists. The total number of peaks dropped for both SMART samples after cleaning (**Fig. 4a**), but the change for 0.1 ng SMART sample was significant larger than that for 1 ng one (**Fig. 4b**), clearly suggesting that with lower amounts of input DNA, more false peaks would be called from the artifact reads (**Fig. 4c**). In support of this, we observed that the 0.1 ng pre-cleaning SMART samples had the largest percentages (on average 64.3%) of peaks located near the poly(T/A) sites (**Fig. 4d**). When compared to the peaks called for the PCR-free data, 51.9% (0.1 ng) and 35.1% (1 ng) of the peaks unique to the pre-cleaning SMART samples overlapped, significantly smaller than the percentages for peaks either shared with or unique to post-cleaned data (**Fig. 4e**). Similarly, the percentages of H3K4me3 peaks (44.4% and 39.8%) located to promoters for the peaks unique to pre-cleaning samples were significantly lower than the numbers for the other two groups of peaks (**Fig. 4f**). As an orthogonal measurement, we analyzed transcription factor (TF) motifs in the H3K4me3 peak regions. The TATA box and CAAT box, two well-known general promoter TF motifs [24], and the ETS motif [25], were the most enriched motifs in the H3K4me3 peaks. In all cases, their occurrences in the peaks detected only in the pre-cleaning samples were significantly lower (**Fig. 4g-i**). In contrast, the RLR1 motif, which basically consists of poly(T), was only enriched in the peaks unique to the pre-cleaning samples (**Fig. 4j**). Finally, we examined the ChIP-seq read densities and aggregated read profiles for the three groups of H3K4me3 peaks, unique to pre- or post-cleaning samples, or shared (**Fig. 4k**). The peaks unique to the post-cleaning samples had about 2x stronger (both 1 ng and 0.1 ng samples) ChIP-seq signals in the PCR-free and Ascel2S data than the peaks unique to the pre-cleaning samples, indicating that the latter peaks were very likely derived from PCR amplification and thus enriched for artifacts (**Fig. 4k**). Taken together, these results indicate that the reads removed by SMARTcleaner are true artifacts and its application can improve the quality of peaks identified from ChIP-seq analysis, resulting in better biological findings.

**Fig. 4.**
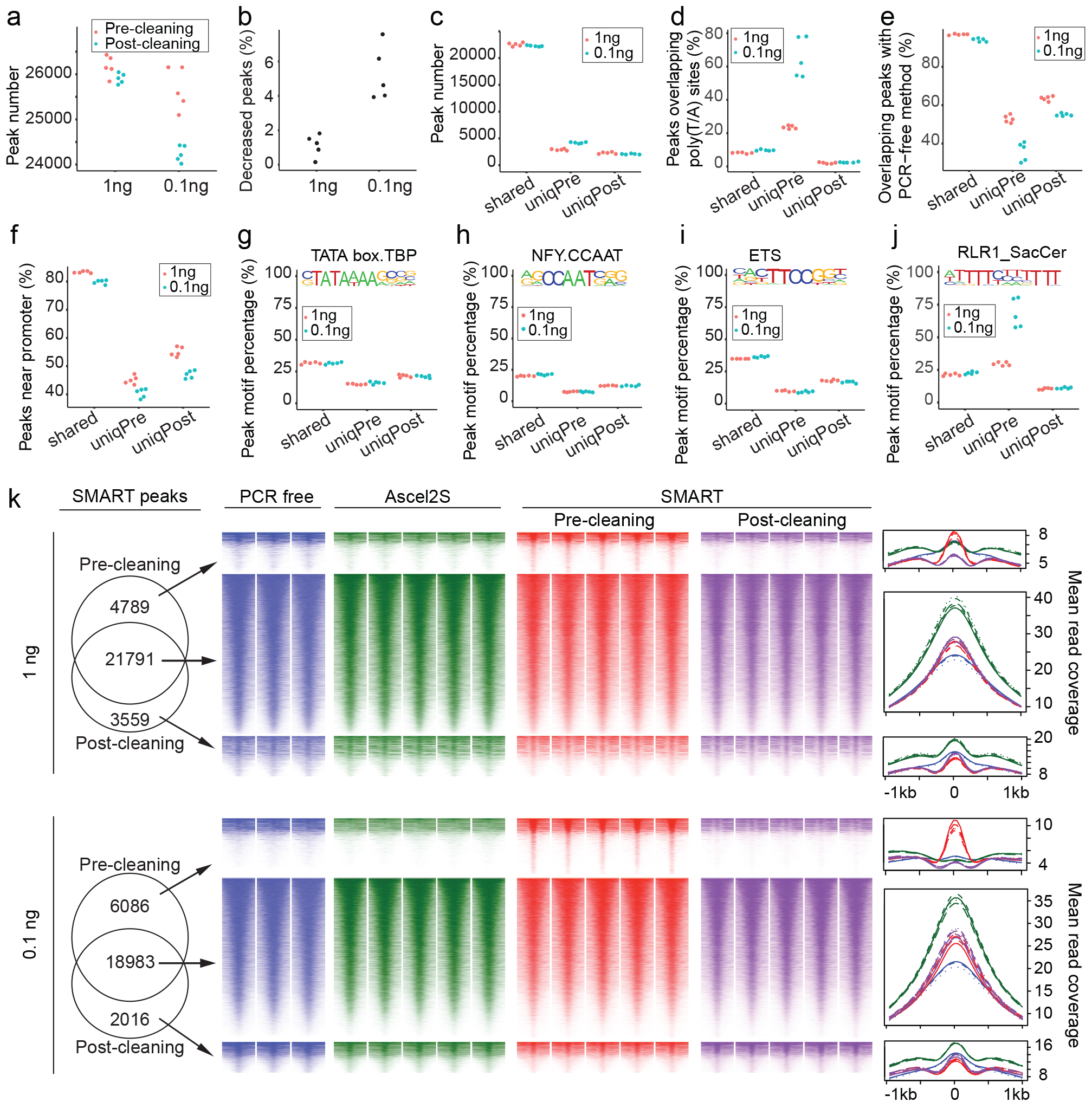
Evaluate SMARTcleaner with H3K4me3 ChIP-seq data. **a**. Numbers of pre- and post-cleaning H3K4me3 peaks. **b**. Change of peak numbers after cleaning. **c**. Numbers of peaks shared or unique to pre-cleaning (“uniqPre”) or post-cleaning (“uniqPost”) data **d**. Overlap of peaks with poly(T/A) sites. **e**. Overlap of peaks with gold standard peaks. **f**. Peaks at the promoter regions (2kb around TSS). **g-j**. Percentages of peaks with each of the four enriched TF motifs. **k**. Read densities and average profiles for peaks shared by or unique to pre- and post-cleaning data. Reads counts were extracted using seqMINER [33] from 6 million reads randomly sampled from individual samples. Heatmaps were drawn using R package pheatmap, with peaks as row and sorted by read densities. In **a-j**, each point represents a replicate sample.

### Evaluation of SMARTcleaner with published transcription factor ChIP-seq datasets

We were especially interested in how the inclusion of artifact reads may affect peaks identified from TF ChIP-seq studies. Therefore, we reanalyzed a previously published Olig2 ChIP-seq dataset (Additional file 1: Table S1, Dataset 3) and compared our results to the original publication [18]. We found that 16% of the original peaks (3,251 of 20,283) overlapped with the poly(T/A) sites, with some peaks exhibiting typical features of false amplification (**Fig. 5a**). We also noticed that the authors applied a combination of very stringent criteria to filter peaks, perhaps in an effort to limit peaks from false priming. Thus, we tried less stringent criteria to obtain a new set of peaks (n=25,179) from the pre-cleaning alignment files and included it in our comparison (see Methods). Next, we used the SMARTcleaner SE mode to clean the alignment files and obtained a list of post-cleaning peaks (n=23,289). A comparison of the three lists of peaks is shown in **Fig. 5b**, from which we defined four groups of peaks (Additional file 2: Figure S4): “TP”, or true positive, called by all methods; “FP”, or false positive, called by the original study and present in the pre-cleaning sample only; “FN”, or false negative, removed by the original study only; and “TN”, or true negative, removed in the original study and by SMARTcleaner. Intersections of the four groups of peaks with poly(T/A) sites showed that 92.9% of TN peaks and 94.3% of FP peaks overlapped with poly(T/A) sites, compared to 12.7% of TP peaks and 5.3% of FN peaks (**Fig. 5c**), indicating that the original study not only included some artifact peaks but also filtered out some true peaks. This was supported by a comparative analysis of the ChIP-seq read intensities, with reads from false priming present in both the ChIP sample and input control (FP and TN in **Fig. 5d,e**). This analysis also showed that the FN group represented true peaks filtered out by the authors by using overly strict criteria (**Fig. 5d,e**).

**Fig. 5.**
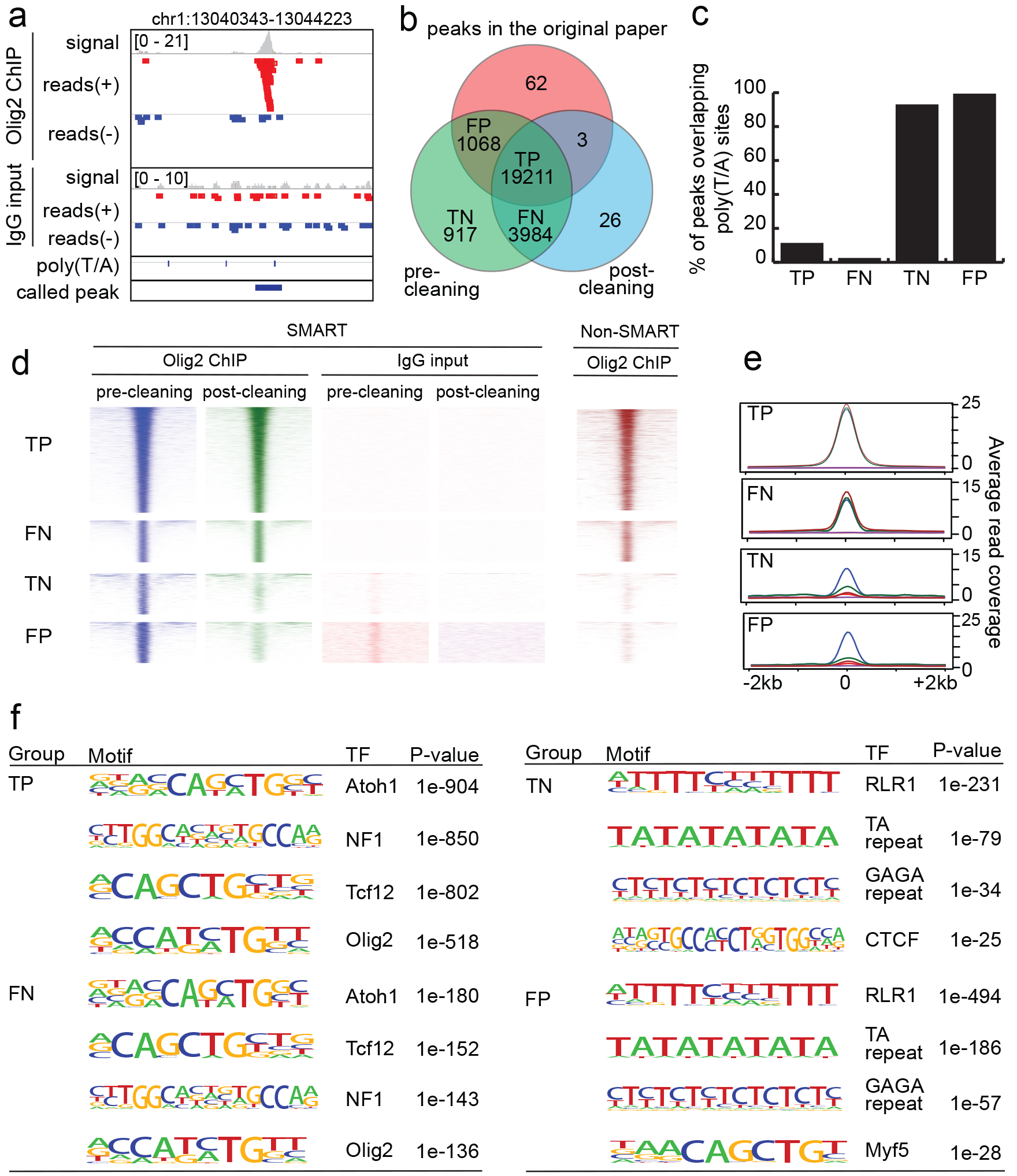
Evaluate SMARTcleaner with a TF ChIP-seq data. **a**. An example of false peaks in the original list of Olig2 ChIP-seq peaks. The track of “called peak” shows peaks provided by the authors. **b**. Venn diagram showing the peak overlaps from three methods: the original peaks from the authors, the peaks called before cleaning, and the peaks called after cleaning. When counting the overlapping peaks, we could get two different numbers depending on which set of peaks is used to report the number (one peak in one set may overlap more than one peak in another set). We reported the smaller number here. **c**. Peaks overlapping with poly(T/A) sites. **d,e**. Read densities and average counts at the four selected groups of peaks, computed by sampling 5 million reads. An Olig2 ChIP-seq data (right) from non-SMART method was also analyzed. **f**. Top enriched motifs by HOMER [30].

To further test the cleaning effect, we included a Olig2 ChIP-seq dataset that was independently generated from neural stem cells using a non-SMART protocol [26]. We found that 86.2% and 91.8% of the pre-cleaning and post-cleaning peaks were detected by the non-SMART method, respectively. Moreover, among the four groups of peaks, 93.8% and 83% of TP and FN peaks were present in the non-SMART peaks, respectively, in contrast to 8.7% and 6.2% for the TN and FP groups, respectively, indicating that false peaks were removed by our clearing process. This result was supported by the patterns in the read density heatmaps and profiles (**Fig. 5d,e**).

In addition, motif analysis demonstrated that the top four motifs enriched in the TP and FN peaks were the same TF motifs (Atoch1, NF1, Tcf12 and Olig2) reported in the original study [18]. However, the top motifs for the TN and FP groups were RLR1, TA repeat, GAGA repeat, CTCF and Myf5, which seem irrelevant to Olig2 function (**Fig. 5f**).

In short, our analysis of the Olig2 ChIP-seq data further supports the value of our newly developed SMARTcleaner tool, and illustrates the need for appropriately removing noise and artifacts from false priming in TF ChIP-seq studies that use the SMART protocol.

### Prevalence of artifact reads from false priming and amplification in SMART-based ChIP-seq datasets

To determine if false priming and amplification is a common problem in SMART-based ChIP-seq libraries, we collected and analyzed all such datasets except a clinical one that is not publicly accessible [15] (Additional file 1: Table S1; see Methods). These ChIP-seq data were carried out in human [13-16], mouse [17, 18], and yeast samples [19]. All but two of the datasets were analyzed by single-end sequencing [14, 15]. Our analysis showed that all available datasets contained an average of 8.5% (2.7% ~19.6%) reads that were likely derived from false priming, regardless of the amount of input DNA (from 0.1 ng to 10 ng DNA) or cell numbers (from 10 to 100 millions) (Additional file 1: Table S1).

## Discussion

The SMART ChIP-seq kit uses the template switching method to improve the efficiency of library construction, which is especially suitable for analyzing samples with very low amounts of input DNA [7]. Consistent with a recent report [14], we show that the protocol, however, can introduce significant noise to ChIP-seq data, due to the annealing of DNA SMART poly(dA) primers to non-targeted genomic regions containing ≥ 12 Ts or As. The artifact reads have distinct features (**Fig. 1**, Additional file 2: Figure S2) that are exploited by the SMARTcleaner tool developed in this study. Using multiple published ChIP-seq datasets, we demonstrated convincingly that our tool can successfully remove the artifact reads arising from false priming and amplification of the SMART poly(dA) primers. It works for both PE and SE ChIP-seq reads (**Fig. 2**, **Fig. 3**), and outputs both cleaned alignment files and noise, which can be loaded into a genome browser for inspecting the cleaning effects visually. SMARTcleaner also provides some running options and helper tools to prepare the files required for the cleaning process. Currently SMARTcleaner does not deal with biases introduced by other factors, such as DNA shearing method etc. [5], but users can easily adapt this tool to their ChIP-seq analytic pipelines and develop it further.

We have examined all currently available public datasets that were obtained using the DNA SMART ChIP-seq kit, and found that the false priming issue is prevalent, regardless of the amount of input DNA material or cell numbers (Additional file 1: Table S1). While the artifact cannot be easily removed by data normalization, strict filtering in peak calling, or a simple exclusion of peaks located at poly(A/T) sites, our study suggests that the false priming issue becomes less severe when a large amount of DNA is used as the starting material for ChIP library preparation. Conceivably, the concern can also be alleviated if high affinity antibodies are used to significantly enrich target DNA templates in the input material. Based on our survey of all available datasets, we have the following recommendations to users of the SMART ChIP-seq kit to exploit its full potential. First, one should use a sufficient amount of DNA as the starting templates, whenever possible. Second, the T-tailing step in the SMART ChIP-seq protocol should be optimized. Third, sequence the NGS libraries using the PE method and clean the ChIP-seq reads using the PE mode of SMARTcleaner. Forth, if the libraries have already been sequenced using the SE method, clean the ChIP-seq reads using the SE mode of SMARTcleaner. Alternatively, one can consider to use other ChIP-seq library preparation methods that can also handle low-input DNA [13, 20, 21].

## Conclusions

False priming and amplification occur at poly(T/A) genomic sites due to the use of poly(dA) primers in SMART-based ChIP-seq library construction. Reads from subsequent false amplification and sequencing are strand-specific and can be effectively removed by our SMARTcleaner tool, leading to improvement in peak calling, and downstream data analysis and interpretation.

## Methods

### ChIP-seq datasets and read processing

The SMART ChIP-seq kit is a promising but relatively new protocol for analyzing small amount of chromatin materials. We searched for ChIP-seq datasets that used this kit in the GEO and by Google and found one publication in 2015 [18], two in 2016 [13, 19], and four in 2017 [14–17]. Among the seven publications, six have made their data publicly accessible (Additional file 1: Table S1). The seventh is a clinical study and the corresponding data have not been released, possibly due to protection of privacy [15]. In the alignment of ChIP-seq reads derived from the SMART protocols, the first three bases were trimmed from the first read (Read1). In all datasets, replicates were analyzed independently. To facilitate comparison with the original studies, we used the same versions of software as in the original publication when applicable.

#### Dataset 1

The first dataset is actually a ChIP-seq of input DNAs from HCT116 cells and HeLa-S3 because the DNA templates were not enriched with any antibodies. It contained seven sets of paired-end sequencing data, which we downloaded from the NCBl SRA database (SRP071830) [14]. Three libraries were constructed using the DNA SMART ChIP-Seq kit (Clontech, #634865), with the others by “standard” ligation-based method. Reads were mapped to the human genome (hg38) using Bowtie2 (v2.2.3) [27], using default parameters with the maximum fragment length for valid paired-end reads set to 2000. Only uniquely mapped reads were kept for further analyses, after duplicate reads were removed using the Picard tool -- MarkDuplicates (v2.3.0, http://broadinstitute.github.io/picard/index.html). To mimic single-end sequencing, we generated SE bam files by extracting the first reads from the PE bam files (samtools view -h -f 64).

#### Dataset 2

The H3K4me3 ChIP experiments were done with 56 million HeLa cells in 56 ChIP reactions [13]. The ChIP DNA was combined into a single pool and then divided into seven aliquots for different library preparation methods and the PCR-free method. Libraries starting from either 1 ng or 0.1 ng ChlP DNA were generated. Reads were aligned to the hg38 human reference genome using Bowtie (v1.2.1) [28]. Only uniquely mapped reads were used for analysis, with duplicate reads removed by samtools (v0.1.19) [29]. To call peaks, we randomly subsampled 6 million mapped reads for each sample, as done in the original study [13] and used the MACS2 (v2.1.0) [23] with *q* value < 0.05. Motif analysis was done using the HOMER (v4.7) [30].

#### Dataset 3

The Olig2 ChIP-seq was carried out with 10 million neural stem cells (NSCs) derived from embryonic (E14.5) CD-1 mice. The libraries were constructed using the DNA SMART ChIP-seq kit and sequenced by the single-end method on an Illumina HiSeq2000 sequencer [18]. The dataset was downloaded from the GEO database (GEO: GSE74646). Reads were aligned to the mouse reference genome (mm10) using bowtie (v1.2.1). Only uniquely mapped reads were used for analysis, with duplicate reads removed using samtools (v0.1.19). Peaks were called using the MACS (v1.4.2) and filtered by *p* value < 10^−5^, fold enrichment > 5, and tag number > 15. When the filter was set to the same as used in the original paper (*p* value < 10^−9^, fold enrichment > 5, and tag number > 20), we obtained essentially the same peaks that were called in the original study. Peak motif analysis was done using HOMER (v4.7) [30].

#### Dataset 4

The H3K4me1 ChIP-seq was obtained with 10 million SUM159 cells. H3K4me1 ChIP-seq libraries were constructed using the DNA SMART ChIP Seq Kit (Clontech) with 10ng ChIP DNA (NCBI GEO: GSE87424) [16]. Raw fastq sequences were downloaded from the GEO and processed with the same methods as the original study.

#### Dataset 5

The ChIP-seq experiments of H3K27ac histone modification and c-MYC were performed with FACS-sorted Eph4 cells. Libraries were constructed using the Clontech DNA Smart Chipseq kit (Clontech, #634866), and pooled for sequencing (NCBI GEO: GSE98004) [17]. Raw fastq sequences were downloaded from the GEO and processed as the original study.

#### Dataset 6

The last dataset was from a yeast study [19]. DNA-RNA immunoprecipitation and deep sequencing (DRIP-seq) was done with S9.6 monoclonal antibody in 100 million yeast cells. We downloaded the alignment files from European Nucleotide Archive (ENA) website (PRJEB8021) and yeast reference genome from the UCSC genome browser [31].

### SMARTcleaner

The SMARTcleaner tool was developed in Perl under the MIT license after analysis of the characteristics of ChIP-seq reads derived from false priming and amplification. Two modes, PE mode and SE mode, were implemented based on the sequencing methods used in ChIP-seq data.

#### PE mode

When sequenced in PE method, the second reads of the falsely primed fragments will pile up upstream of the poly(T) sites or the downstream of the poly(A) sites (**Figure 1e,f**), allowing two mismatch insertions (Additional file 2: Figure S2). SMARTcleaner will go through a sorted (by coordinates) alignment file and find read pairs with the second read at the left end of poly(T) sites or at the right end of poly(A) sites (Additional file 2: Figure S5). It will keep tracking the number of such fragments at each position of a poly(T/A) site. When this number is over a threshold (default: 1) predefined for false amplification, all read pairs ending in the same position will be considered as artifacts and placed to the new alignment file (“noise bam file”). In the meantime, the original bam file subtracting the artifact reads will be saved as a cleaned bam file.

#### SE mode

When ChIP-seq is sequenced in SE method, the false reads will be clustered upstream of poly(T) sites or downstream of poly(A) sites of the reference genome (**Fig. 1**), up to two mismatches (Additional file 2: Figure S2). SMARTcleaner first examines the reads in the flanking regions (by default 2kb) of all poly(T/A) sites to decide the size of the region containing falsely amplified fragments. For reads on “+” strand, the distance is calculated from the left ends of reads to the left ends of poly(T) sites (Additional file 2: Figure S6a) or the right ends of poly(A) sites (Additional file 2: Figure S6b). For reads on “−” strand, the distance is calculated from the right ends of reads to the left ends of poly(T) sites (Additional file 2: Figure S6a) or the right ends of poly(A) sites (Additional file 2: Figure S6b). Based on the distribution of the distances, SMARTcleaner automatically determines the window size at poly(T/A) sites for sampling, or a user can manually set it according to the read distribution at the poly(T/A) sites (**Fig. 1h,i**). A bed file containing the resampling regions will be generated. Next, it will go through the reads at each of those regions, check if the potentially artifact reads outnumber (default 2x) those in the unaffected opposite strand, and finally resample the artifact reads, if necessary, according to the read numbers in the opposite strand (Additional file 2: Figure S7a,b). For the genomic regions with overlapping poly(T) and poly(A) sites, the tool will process the poly(T/A) sites based on the order of their appearance in the reference genome (Additional file 2: Figure S7c).

For SE mode, a list of poly(T/A) sites is needed. We included a helper command to identify such regions in a genome. To estimate the range for resampling reads, we implemented another helper command in our tool for this purpose. Users can also directly set a range for resampling based on their knowledge of their datasets or the fragment distribution around the poly(T/A) sites.

## List of abbreviations

ChIP-seq: chromatin immunoprecipitation and deep sequencing
NGS: next-generation sequencing
PE: paired-end
SE: single-end
SMART: switching mechanism at the 5’ end of the RNA transcript

## Declarations

### Ethics approval and consent to participate

Not applicable

### Consent for publication

Not applicable

### Availability of data and material

The datasets supporting the conclusions of this article are available in the NCBI SRA: SRP071830 (Dataset 1), NCBI SRA: SRP067250 (Dataset 2), NCBI GEO: GSE74646 (Dataset 3), NCBI GEO: GSE87424 (Dataset 4), NCBI GEO: GSE98004 (Dataset 5), and European Nucleotide Archive (ENA): PRJEB8021 (Dataset 6). SMARTcleaner is publicly available under MIT license at github (https://github.com/dzhaobio/SMARTcleaner).

### Authors’ contributions

D. Zhao developed the tool, performed the analyses, and drafted the manuscript. D. Zheng contributed ideas, wrote the manuscript, and supervised the study. All authors read and approved the final manuscript.

## Acknowledgements

We thank the High Performance Computing Core of Albert Einstein College of Medicine for their computational support. We thank Dr. Itamar Simon for sharing additional information about the dataset 1. We also would like to thank Yilin Zhao, Ping Wang, and Yang Liu for helpful discussions, Feng Jiang, Pengcheng Yang and Mingyan Lin for testing the SMARTcleaner tool, and Herbert Lachman for manuscript editing.

## Description of additional data files

Additional file 1: Supplementary Table S1.

Additional file 2: Supplementary Figure S1–S7.

